# Mitochondrial citrate transporters CtpA and YhmA are involved in lysine biosynthesis in the white koji fungus, *Aspergillus luchuensis* mut*. kawachii*

**DOI:** 10.1101/341370

**Authors:** Chihiro Kadooka, Kosuke Izumitsu, Masahira Onoue, Kayu Okutsu, Yumiko Yoshizaki, Kazunori Takamine, Masatoshi Goto, Hisanori Tamaki, Taiki Futagami

## Abstract

*Aspergillus luchuensis* mut*. kawachii* produces a large amount of citric acid during the process of fermenting shochu, a traditional Japanese distilled spirit. In this study, we characterized *A. kawachii* CtpA and YhmA, which are homologous to the yeast *Saccharomyces cerevisiae* mitochondrial citrate transporters Ctp1 and Yhm2, respectively. CtpA and YhmA were purified from *A. kawachii* and reconstituted into liposomes. The proteoliposomes exhibited only counter-exchange transport activity; CtpA transported citrate using counter substrates especially for *cis-*aconitate and malate, whereas YhmA transported citrate using a wider variety of counter substrates, including citrate, 2-oxoglutarate, malate, *cis*-aconitate, and succinate. Disruption of *ctpA* and *yhmA* caused deficient hyphal growth and conidia formation with reduced mycelial weight–normalized citrate production. Because we could not obtain a Δ*ctpA* Δ*yhmA* strain, we constructed a *ctpA-S* conditional expression strain in the Δ*yhmA* background using the Tet-On promoter system. Knockdown of *ctpA*-*S* in Δ*yhmA* resulted in a severe growth defect on minimal medium, indicating that double disruption of *ctpA* and *yhmA* leads to synthetic lethality; however, we subsequently found that the severe growth defect was relieved by addition of lysine. Our results indicate that CtpA and YhmA are mitochondrial citrate transporters involved in citric acid production and that transport of citrate from mitochondria to the cytosol plays an important role in lysine biogenesis in *A. kawachii*.

**IMPORTANCE:** Citrate transport is believed to play a significant role in citrate production by filamentous fungi; however, details of the process remain unclear. This study characterized two citrate transporters from *Aspergillus luchuensis* mut*. kawachii*. Biochemical and gene disruption analyses showed that CtpA and YhmA are mitochondrial citrate transporters required for normal hyphal growth, conidia formation, and citric acid production. In addition, this study provided insights into the links between citrate transport and lysine biosynthesis. The characteristics of fungal citrate transporters elucidated in this study will help expand our understanding of the citrate production mechanism and facilitate the development and optimization of industrial organic acid fermentation processes.

## INTRODUCTION

The white koji fungus, *Aspergillus luchuensis* mut. *kawachii* (*A. kawachii*), is a filamentous fungus used for the production of shochu, a traditional Japanese distilled spirit (1, 2). During the shochu fermentation process, *A. kawachii* secretes large amounts of the glycoside hydrolases α-amylase and glucoamylase, which degrade starches contained in cereal ingredients such as rice, barley, and sweet potato (3). The resulting monosaccharides or disaccharides can be further utilized by the yeast *Saccharomyces cerevisiae* for ethanol fermentation. In addition to this feature, *A. kawachii* produces a large amount of citric acid, which lowers the pH of the “*moromi*” (mash) to between 3 and 3.5, thereby preventing the growth of contaminant microbes. This feature is important because shochu is mainly produced in relatively warm areas of Japan, such as Kyushu and Okinawa islands.

Although a clearly different species, *A. kawachii* is phylogenetically closely related to *Aspergillus niger*, which is commonly used in the citric acid fermentation industry (4-6). The mechanism of citric acid production by *A. niger* has been investigated from various perspectives, and related metabolic pathways have been elucidated (7-9). Carbon sources such as glucose and sucrose are metabolized to produce pyruvate via the glycolytic pathway; subsequently, citric acid is synthesized by citrate synthase as an intermediate compound of the tricarboxylic acid cycle in mitochondria and excreted to the cytosol prior to subsequent excretion to the extracellular environment. A previous study detected citrate synthase activity primarily in the mitochondrial fraction (10). Experiments involving overexpression of the citrate synthase–encoding *citA* gene indicated that citrate synthase plays only a minor role in controlling the flux of the pathway involved in citric acid production (11). By contrast, a mathematical analysis suggested that citric acid overflow might be controlled by the transport process (e.g., uptake of carbon source, pyruvate transport from the cytosol to mitochondria, transport of citrate from mitochondria to the cytosol and then extracellular excretion) (12-14).

Mitochondrial citrate transporters of mammals and *S. cerevisiae* have been well characterized. Biochemical studies revealed that rat liver citrate transporter (CTP) catalyzes the antiport reaction of the dibasic form of tricarboxylic acids (e.g., citrate, isocitrate, and *cis-*aconitate) with other tricarboxylic acids, dicarboxylic acids (e.g., malate, succinate, maleate), or phosphoenolpyruvate (15-17), whereas the *S. cerevisiae* citrate transporter (Ctp1) shows stricter substrate specificity for tricarboxylic acids compared with CTP (18, 19). Cytosolic citrate is used in the production of acetyl-CoA, which plays a significant role in the biosynthesis of fatty acids and sterols in mammalian cells (20-23). However, no phenotypic changes were observed in *S. cerevisiae* following disruption of the *CTP1* gene (24), perhaps because other transport processes (e.g., involving mitochondrial succinate/fumarate transporter Acr1) control acetyl-CoA synthesis (25-27). A homolog to the *CTP1* gene (*ctpA*) was recently characterized in *A. niger*, with a focus on the relationship between citrate transport and the organism’s high citrate production capability (28). Disruption of the *ctpA* gene led to reduced growth and citric acid production in *A. niger* only during the early logarithmic phase, indicating that CtpA is not a major mitochondrial citrate transporter in *A. niger* (28).

To better understand the mechanism of citric acid production by *A. kawachii*, we previously characterized the changes in gene expression that occur during solid-state culture, which is used for brewing shochu (29). During the shochu-making process, the cultivation temperature is tightly controlled, with gradual increase to 40°C and then lowering to 30°C. Lowering of the temperature is required to enhance production of citric acid (30). We sought to identify genes related to citric acid production and reported that expression of the gene encoding the putative mitochondrial citrate/malate transporter (AKAW_03754, CtpA) increased 1.78-fold upon lowering of the temperature (29). Subsequently, we found that expression of a putative mitochondrial citrate transporter (AKAW_06280, YhmA) gene, which is a homolog of the mitochondrial citrate/2-oxoglutarate transporter Yhm2 of *S. cerevisiae* (31), increased 1.76-fold, based on analysis of a microarray dataset (29). Yhm2 of *S. cerevisiae* was first characterized as a DNA-binding protein predicted to play a role in replication and segregation of the mitochondrial genome (32). However, subsequent biochemical and genetic studies revealed that Yhm2 is a mitochondrial transporter that catalyzes the antiport reaction of citrate and 2-oxoglutarate (31). Yhm2 also exhibited transporter activity for oxaloacetate, succinate, and to a lesser extent, fumarate. Yhm2 plays a significant physiologic role in the citrate/2-oxoglutarate NADPH redox shuttle in *S. cerevisiae* to reduce levels of reactive oxygen species.

In this study, we focused on characterizing both CtpA and YhmA of *A. kawachii* to uncover the functional role of these putative mitochondrial citrate transporters. Our biochemical analyses of purified CtpA and YhmA and phenotypic analyses of disruptant strains confirmed that CtpA and YhmA are mitochondrial citrate carriers involved in citric acid production. Our findings also suggest that double disruption of *ctpA* and *yhmA* induces a synthetic lethal phenotype in minimal (M) medium and that the process of transporting citrate from mitochondria to the cytosol is of physiologic significance for lysine biosynthesis in *A. kawachii*.

## RESULTS

### Sequence features of CtpA and YhmA

The *A. kawachii* genes *ctpA* and *yhmA* encode proteins of 296 and 299 amino acid residues, respectively. These proteins were found to contain six predicted transmembrane domains and three P-X-(D/E)-X-X-(R/K) sequences, which are common characteristics of mitochondrial carrier proteins (33-35) (Fig. S1 in the supplemental material).

Amino acid sequence identities of 47, 35, and 71% were determined between *A. kawachii* CtpA and *S. cerevisiae* Ctp1, between *A. kawachii* CtpA and rat CTP, and between *A. kawachii* YhmA and *S. cerevisiae* Yhm2, respectively. The amino acid residues required for interacting with citrate in *S. cerevisiae* Ctp1 (site I [K83, R87, and R189] and site II [K37, R181, K239, R276, and R279]) were conserved in *A. kawachii* CtpA (36, 37) (Fig. S1A in the supplemental material). The predicted substrate binding sites in *S. cerevisiae* Yhm2 (site I [E83, K87, and L91], site II [R181 and Q182], and site III [R279]) were also conserved in *A. kawachii* YhmA (31, 35) (Fig. S1B in the supplemental material).

The *A. kawachii* genome contained an additional *S. cerevisiae yhm2* homologous gene, *yhmB* (AKAW_02589) (Fig. S1B in the supplemental material), which encodes a protein of 309 amino acid resides with a 53% sequence identity to Yhm2. All of the amino acid residues of sites I, II, and III in Yhm2 mentioned above were found to be conserved in YhmB, suggesting that YhmB functions as a mitochondrial carrier protein. However, no *yhmB* transcripts were detected in microarray analyses during the shochu fermentation process (29). In addition, disruption of *yhmB* did not induce a phenotypic change in *A. kawachii* in M medium (data not shown). Thus, we excluded analysis of the *yhmB* gene from this study.

### Transport activity of CtpA and YhmA

To clarify whether CtpA and YhmA are citrate transporters, we purified the proteins and assayed their activity. For the purification of CtpA and YhmA, C-terminal S-tag fusion proteins were expressed in *A. kawachii* Δ*ctpA* and Δ*yhmA* strains, respectively, under control of the Tet-On promoter, and the products were purified using S-protein agarose (Fig. S2 in the supplemental material). Purified CtpA and YhmA were reconstituted into liposomes using a freeze/thaw sonication procedure, and then [^14^C]-citrate uptake into proteoliposomes was assessed as either uniport (absence of internal substrate) or antiport (presence of internal substrate such as oxaloacetate, succinate, *cis*-aconitate, citrate, 2-oxoglutarate, or malate). Uptake of [^14^C]-citrate was only observed under antiport conditions for both CtpA and YhmA reconstituted proteoliposomes (Fig. 1A and B). CtpA exhibited higher specificity for *cis-*aconitate and malate and showed activity in the presence of oxaloacetate, succinate, and citrate, although to a lesser extent (Fig. 1A). Much lower activity of CtpA was also detected in the presence of 2-oxoglutarate. By contrast, YhmA exhibited wider specificity, with activity toward citrate, 2-oxoglutarate, malate, *cis*-aconitate, and succinate to the same extent and activity in the presence of oxaloacetate to a lesser extent (Fig. 1B).

**Figure 1.**
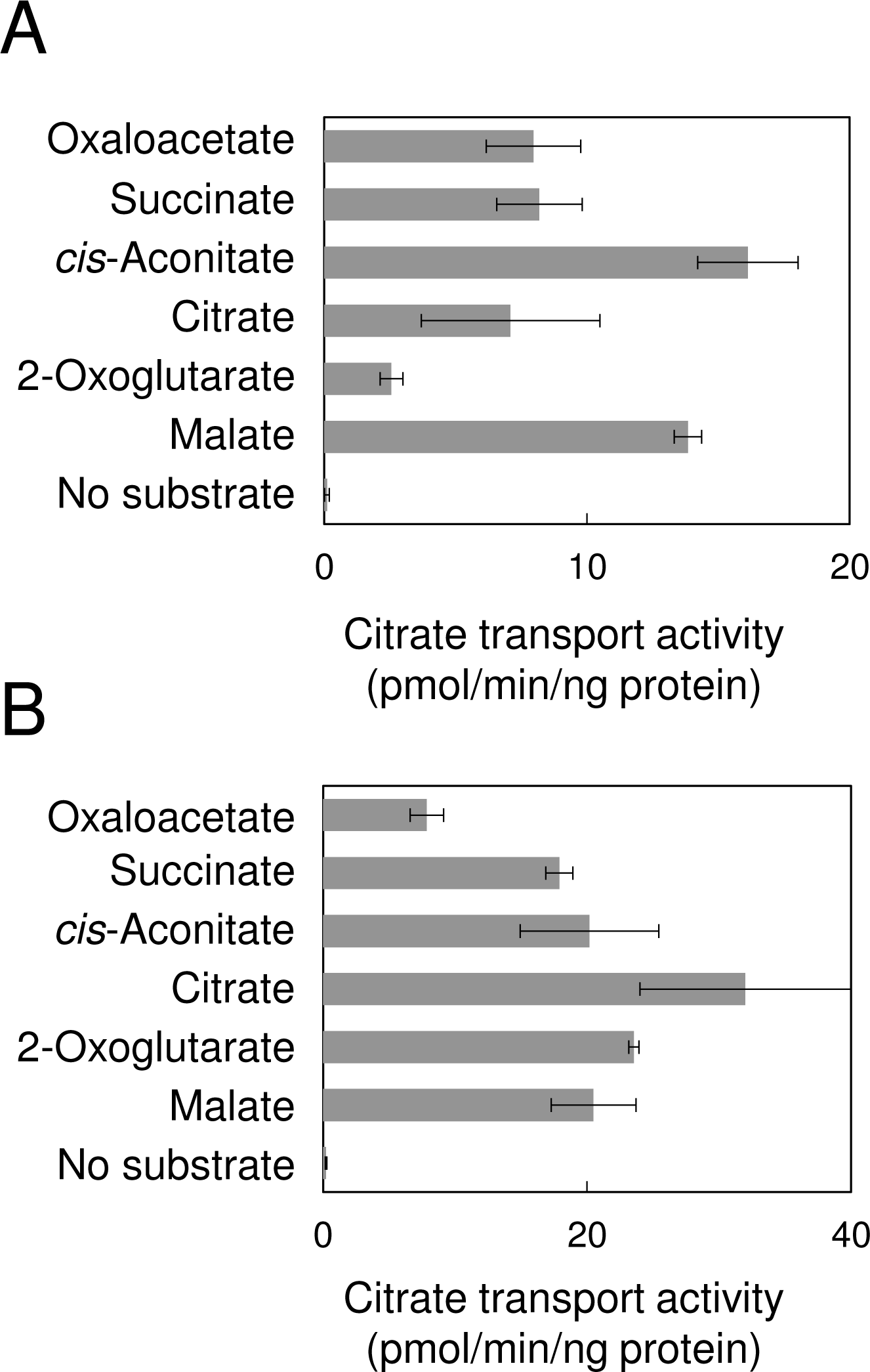
Citrate transport activity of (A) CtpA-S and (B) YhmA-S. CtpA-S or YhmA-S reconstituted proteoliposomes were preloaded with or without 1 mM internal substrate (oxaloacetate, succinate, *cis*-aconitate, citrate, 2-oxoglutarate, or malate). The exchange assay was initiated by adding 1 mM [^14^C]-citrate (18.5 kBq) to the exterior of the proteoliposomes and terminated after 30 min. The mean and standard deviation were determined from the results of 3 independent measurements.

### Phenotype of control, Δ*ctpA*, Δ*yhmA*, and Ptet-*ctpA-S* Δ*yhmA* strains

To explore the physiologic roles of CtpA and YhmA, we characterized the colony morphology of the *A. kawachii* Δ*ctpA* and Δ*yhmA* strains. The Δ*ctpA* strain showed a growth defect at 25 and 30°C, and the defective phenotype was restored at 37 and 42°C (Fig. 2A). This result agreed with a previous report indicating that the *A. niger* Δ*ctpA* strain is more sensitive to low-temperature stress (28). By contrast, the Δ*yhmA* strain exhibited smaller colony diameter than the control strain on M medium at all temperatures tested.

**Figure 2.**
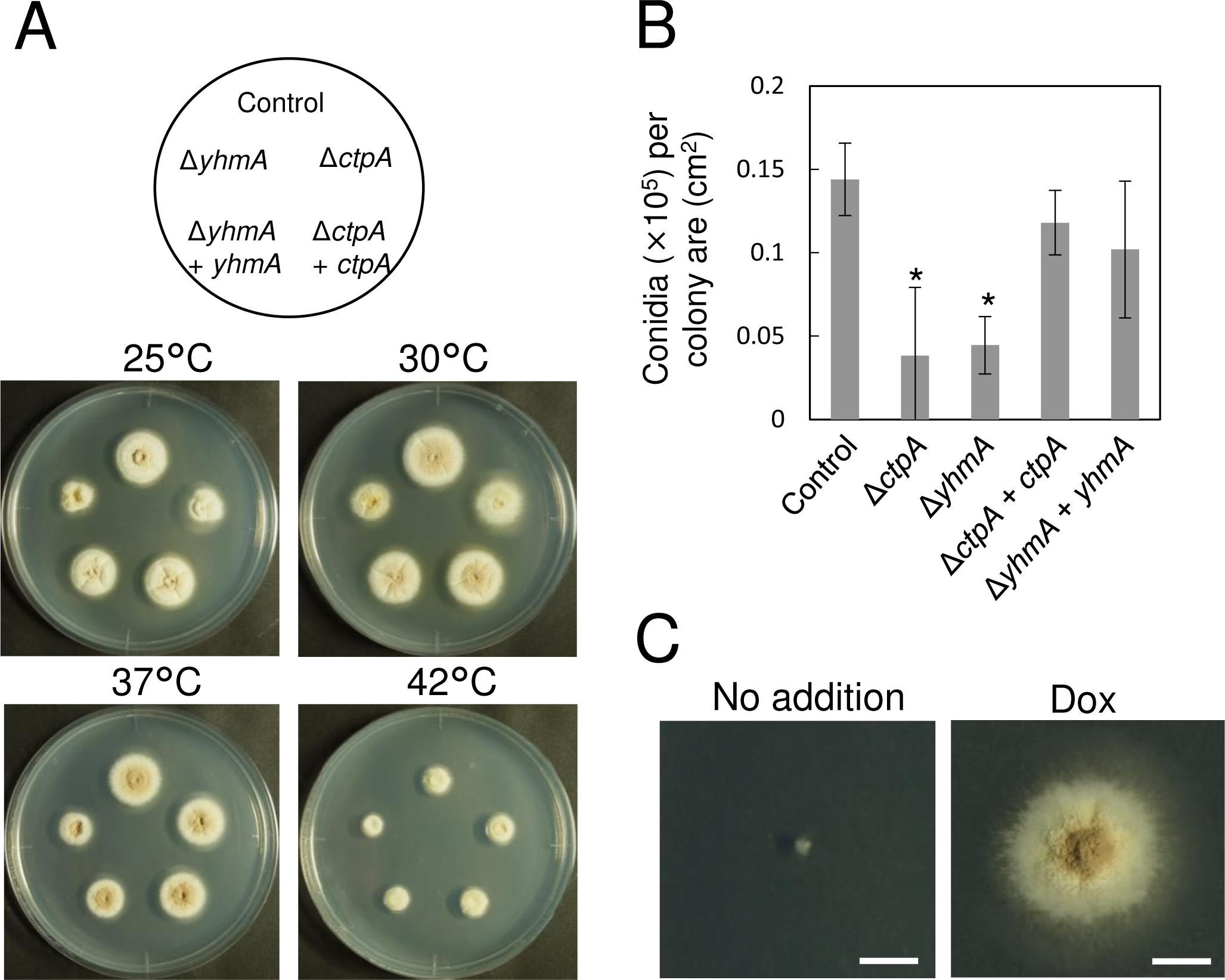
(A) Morphology of *A. kawachii* colonies. Conidia (10^4^) were inoculated onto M agar medium and incubated for 4 days. (B) Conidia formation on M agar medium. Conidia (10^4^) were inoculated onto M agar medium. After 5 days of incubation at 30° C, newly formed conidia were suspended in 0.01% (wt/vol) Tween 20 solution and counted using a hemocytometer. The mean and standard deviation of the number of conidia formed were determined from the results of 3 independently prepared agar plates. *, Statistically significant difference (*p* < 0.05, Welch’s *t*-test) relative to the result for the control strain. (C) Colony formation of the *A. kawachii* Ptet-*ctpA-S* Δ*yhmA* strain. Conidia (10^4^) were inoculated onto M agar medium with or without 1 μg/ml Dox and incubated at 30° C for 5 days. Scale bars indicate 1 cm.

Because the colonies of the Δ*ctpA* and Δ*yhmA* strains were paler in color than colonies of the control strain, we assessed conidia formation (Fig. 2B). Strains were cultivated on M medium at 30°C for 4 days, at which time the number of conidia formed was determined. The number of conidia per cm^2^ of the Δ*ctpA* and Δ*yhmA* strains declined significantly, to approximately 30% of the number produced by the control strain (Fig. 2B), indicating that CtpA and YhmA are involved in conidia formation. Complementation of *ctpA* (Δ*ctpA* + *ctpA*) and *yhmA* (Δ*yhmA + yhmA*) successfully reversed the above-mentioned deficient phenotypes of the Δ*ctpA* and Δ*yhmA* strains.

We then attempted to construct a *ctpA*/*yhmA* double disruptant by disrupting the *yhmA* gene in the Δ*ctpA* strain. However, all of transformants obtained were heterokaryotic gene disruptants (data not shown). Therefore, we constructed a strain that conditionally expressed the S-tagged *ctpA* gene (*ctpA-S*) using the Tet-On system and then disrupted the *yhmA* gene under *ctpA-S*–expressing conditions using doxycycline (Dox), yielding strain Ptet-*ctpA-S* Δ*yhmA*. Dox-controlled expression of CtpA-S was confirmed at the protein level by immunoblot analysis using anti–S-tag antibody (Fig. S3 in the supplemental material, right panel). The Ptet-*ctpA-S* Δ*yhmA* strain exhibited a severe growth defect in M medium without Dox (*ctpA-S* expression is not induced in the absence of Dox) (Fig. 2C), indicating that double disruption of *ctpA* and *yhmA* induces synthetic lethality in M medium.

### Organic acid production by control, Δ*ctpA*, Δ*yhmA*, and Ptet-*ctpA*-*S* Δ*yhmA* strains

To investigate the physiologic role of CtpA and YhmA in organic acid production, we compared organic acid production by the control, Δ*ctpA*, Δ*yhmA*, and Ptet-*ctpA-S* Δ*yhmA* strains (Fig. 3). The control, Δ*ctpA*, and Δ*yhmA* strains were pre-cultivated in M medium at 30°C for 36 h and then transferred to CAP medium and further cultivated at 30°C for 48 h. By contrast, the Ptet-*ctpA-S* Δ*yhmA* strain was pre-cultured in M medium with Dox before transfer to CAP medium without Dox because this strain cannot grow in the absence of Dox (non-induced *ctpA*-*S* expression condition). CAP medium was used for organic acid production because it contains a high concentration of carbon source (10% [wt/vol] glucose) and appropriate trace elements (7-9). We measured organic acid levels in the culture supernatant and mycelia separately as the extracellular and intracellular fractions, respectively.

**Figure 3.**
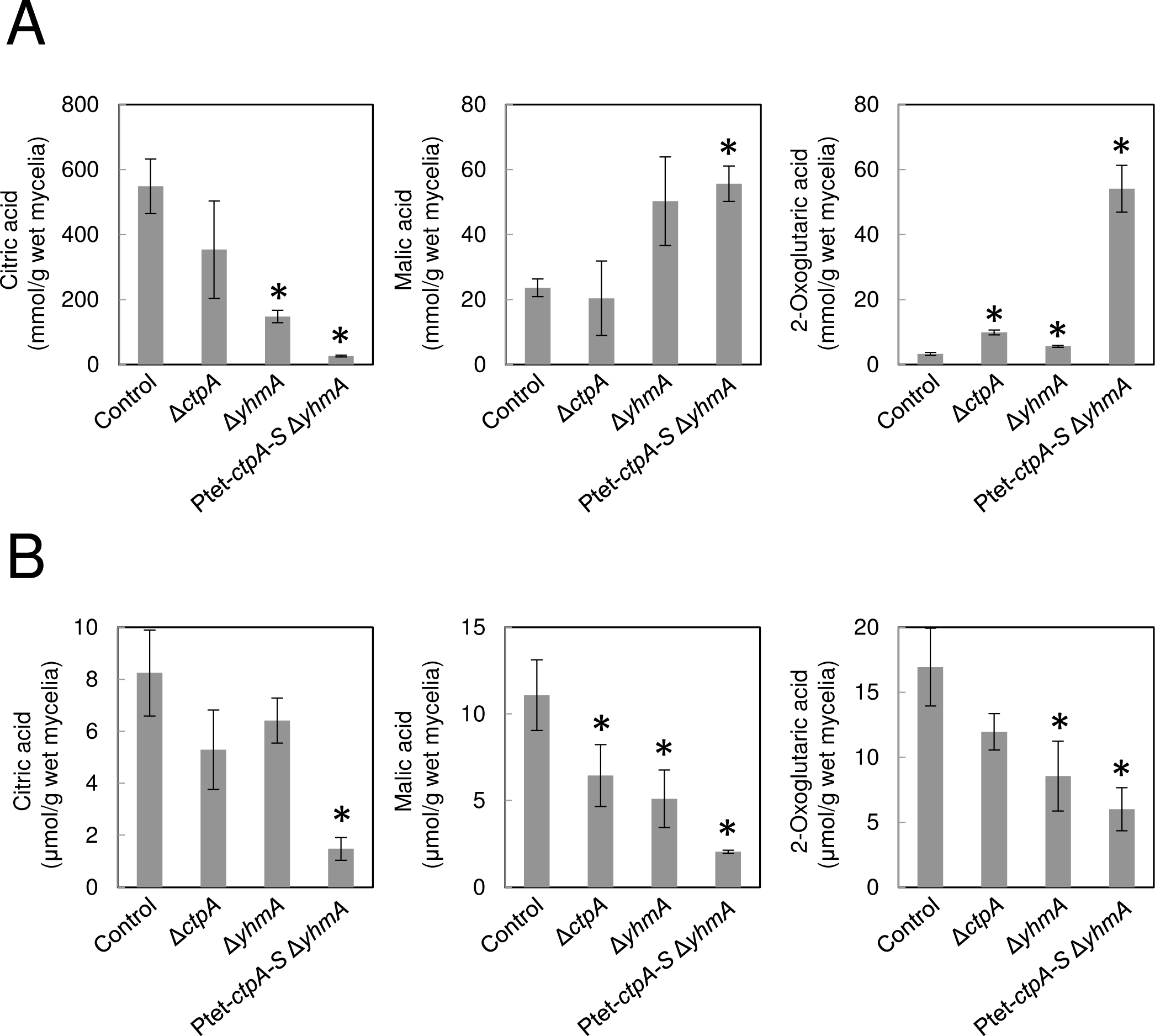
(A) Extracellular and (B) intracellular organic acid production by *A. kawachii* strains. The control, Δ*ctpA*, Δ*yhmA*, and Ptet-*ctpA-S* Δ*yhmA* strains were pre-cultured in M medium for 36 h, then transferred to CAP medium and further cultivated for 48 h. The mean and standard deviation were determined from the results of 3 independent cultivations. *, Statistically significant difference (*p* < 0.05, Welch’s *t*-test) relative to the result for the control strain.

In the extracellular fraction, citric acid was detected as the major organic acid, but malic acid and 2-oxoglutaric acid were also detected using our HPLC system (Fig. 3A). The Δ*ctpA* strain exhibited 3.0-fold greater production of extracellular 2-oxoglutaric acid compared with the control strain, whereas the Δ*yhmA* strain exhibited 0.27-fold lower citric acid and 1.7-fold greater 2-oxoglutaric acid production. In addition, the Ptet-*ctpA*-*S* Δ*yhmA* strain exhibited 0.047-fold lower citric acid production, 2.4-fold greater malic acid production, and 16-fold greater 2-oxoglutaric acid production in the extracellular fraction.

In the intracellular fraction, citric acid, malic acid, and 2-oxoglutaric acid were detected at similar concentrations (Fig. 3B). The decrease in production of citric acid by the Δ*ctpA* and Δ*yhmA* strains was not statistically significant, but the Δ*ctpA* strain exhibited 0.58-fold lower malic acid production, and the Δ*yhmA* strain exhibited 0.46- and 0.50-fold lower malic acid and 2-oxoglutaric acid production, respectively, compared with the control strain. By contrast, the intracellular concentrations of citric acid, malic acid, and 2-oxoglutaric acid produced by the Ptet-*ctpA-S* Δ*yhmA* strain were 0.18-, 0.18-, and 0.35-fold lower than the control, respectively.

These results indicate that CtpA and YhmA play a significant role in organic acid production in *A. kawachii*. The concentration of citric acid produced tended to be negatively correlated with the concentrations of malic acid and 2-oxogluataric acid in the extracellular fraction. In addition, the Ptet-*ctpA-S* Δ*yhmA* strain exhibited the most significant change, especially with regard to the reduced concentration of citric acid in both the extracellular and intracellular fractions, suggesting that CtpA and YhmA function redundantly in citric acid production.

### Transcriptional analysis of *ctpA* and *yhmA*

To investigate the effect of growth phase on expression of the *ctpA* and *yhmA* genes, we performed real-time reverse transcription PCR analysis using RNA extracted from mycelia and conidia of the *A. kawachii* control strain. The control strain was cultivated in M liquid medium or M agar medium for generation of mycelia or conidia, respectively. The growth phase during liquid cultivation was evaluated by measuring the weight of freeze dried mycelia (Fig. 4A). Based on mycelial weight, 0 to 24 h, 24 to 30 h, 30 to 36 h, and 36 to 60 h corresponded to the lag, early log, late log, and stationary phases, respectively. We tested the quality of conidial RNA by assessing production of *wetA* transcripts, which are abundant in dormant *A. niger* conidia (38). The level of *wetA* transcription was 13-fold higher in conidia than mycelia for culture in M liquid medium for 36 h, indicating that extraction of RNA from the conidia was successful (Fig. 4B).

**Figure 4.**
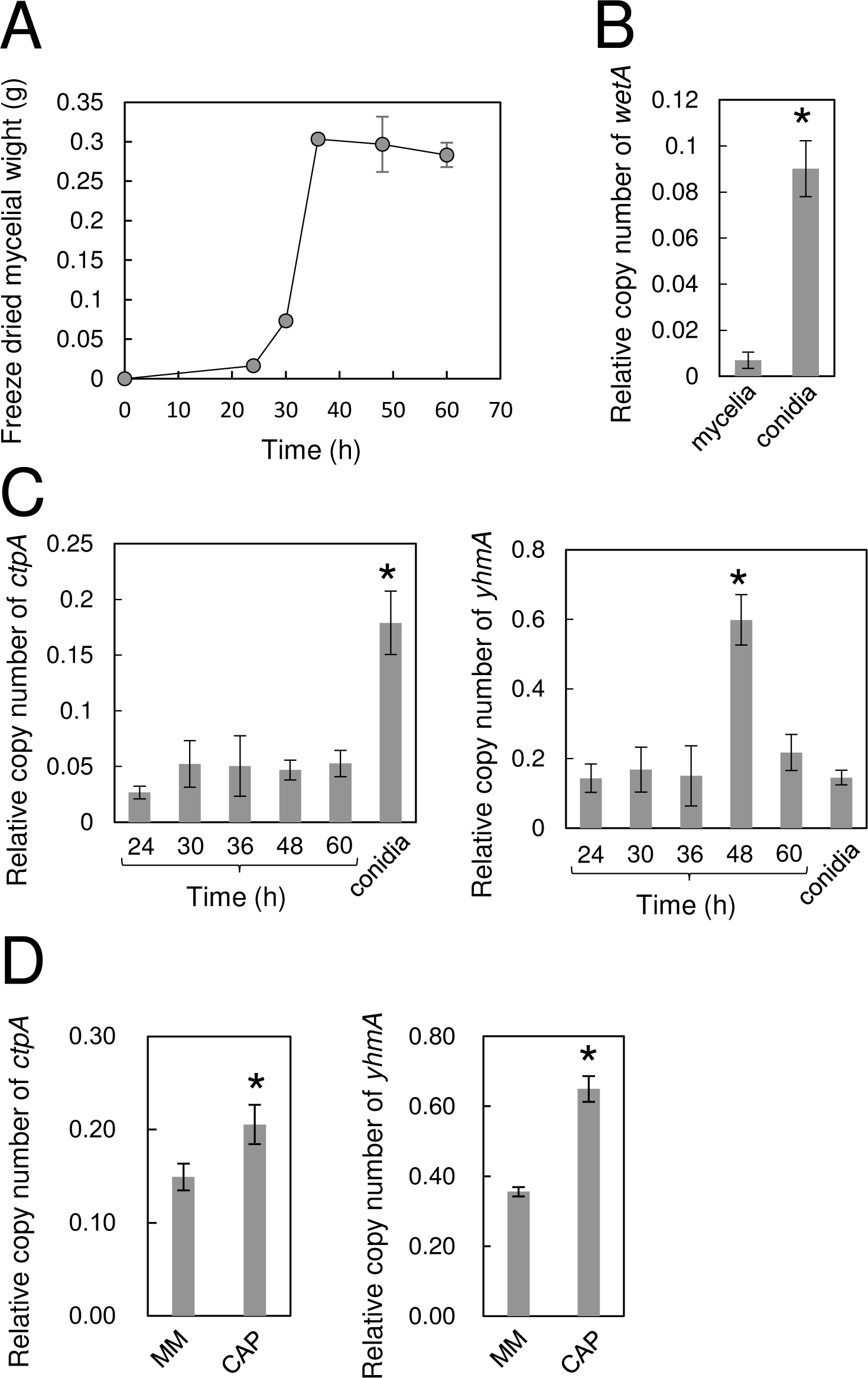
(A) Growth curve of *A. kawachii* in M liquid medium at 30° C. (B) Comparison of relative expression level of *wetA* in mycelia (stationary phase at 36 h) and conidia. (C) Comparison of relative expression levels of *ctpA* and *yhmA* in mycelia and conidia. (D) Comparison of relative expression levels of *ctpA* and *yhmA* in M medium and CAP medium. All results were normalized to the expression level of the actin-encoding gene, *actA*. The mean and standard deviation were determined from the results of 3 independent cultivations. *, Statistically significant difference (*p* < 0.05, Welch’s *t*-test) relative to results obtained under other conditions.

We then investigated transcription of the *ctpA* and *yhmA* genes (Fig. 4C). The level of *ctpA* expression was relatively constant across the vegetative growth period, whereas expression increased significantly in conidia. By contrast, expression of *yhmA* increased significantly at 48 h (stationary phase) and then decreased at 60 h. The level of *yhmA* transcripts in the conidia was similar to that in vegetative hyphae in the lag and log phases.

We also investigated the effect of medium composition (M or CAP medium) on the transcription of *ctpA* and *yhmA* because *A. kawachii* produces a large amount of citric acid in CAP medium but not M medium (Fig. 4D). Expression levels of both *ctpA* and *yhmA* were higher in CAP medium than M medium.

### Subcellular localization of CtpA and YhmA

To determine the subcellular localization of CtpA and YhmA, green fluorescent protein (GFP) was fused to the C-terminus of CtpA and YhmA and expressed in the Δ*ctpA* and Δ*yhmA* strains, respectively, under control of the respective native promoters. Functional expression of CtpA-GFP and YhmA-GFP was confirmed by complementation of the deficient phenotype of the Δ*ctpA* and Δ*yhmA* strains (Fig. 5A). We first examined the strains expressing CtpA-GFP and YhmA-GFP when grown in M medium. Green fluorescence associated with YhmA-GFP merged well with the red fluorescence of MitoTracker Red CMXRos, which stains mitochondria (Fig. 5B). No green fluorescence was detected for the strain expressing CtpA-GFP in M medium, however (data not shown). Because the *ctpA* and *yhmA* genes were transcribed at higher levels in CAP medium than M medium (Fig. 4D), we then cultivated the strains in CAP medium. Green fluorescence associated with YhmA-GFP (Fig. 5C, left panel) and CtpA-GFP (Fig. 5C right panel) was detected, and this fluorescence merged with the red fluorescence, although not completely. This result suggests that CtpA-GFP and YhmA-GFP are localized in the mitochondria, but this should be confirmed through additional experiments.

**Figure 5.**
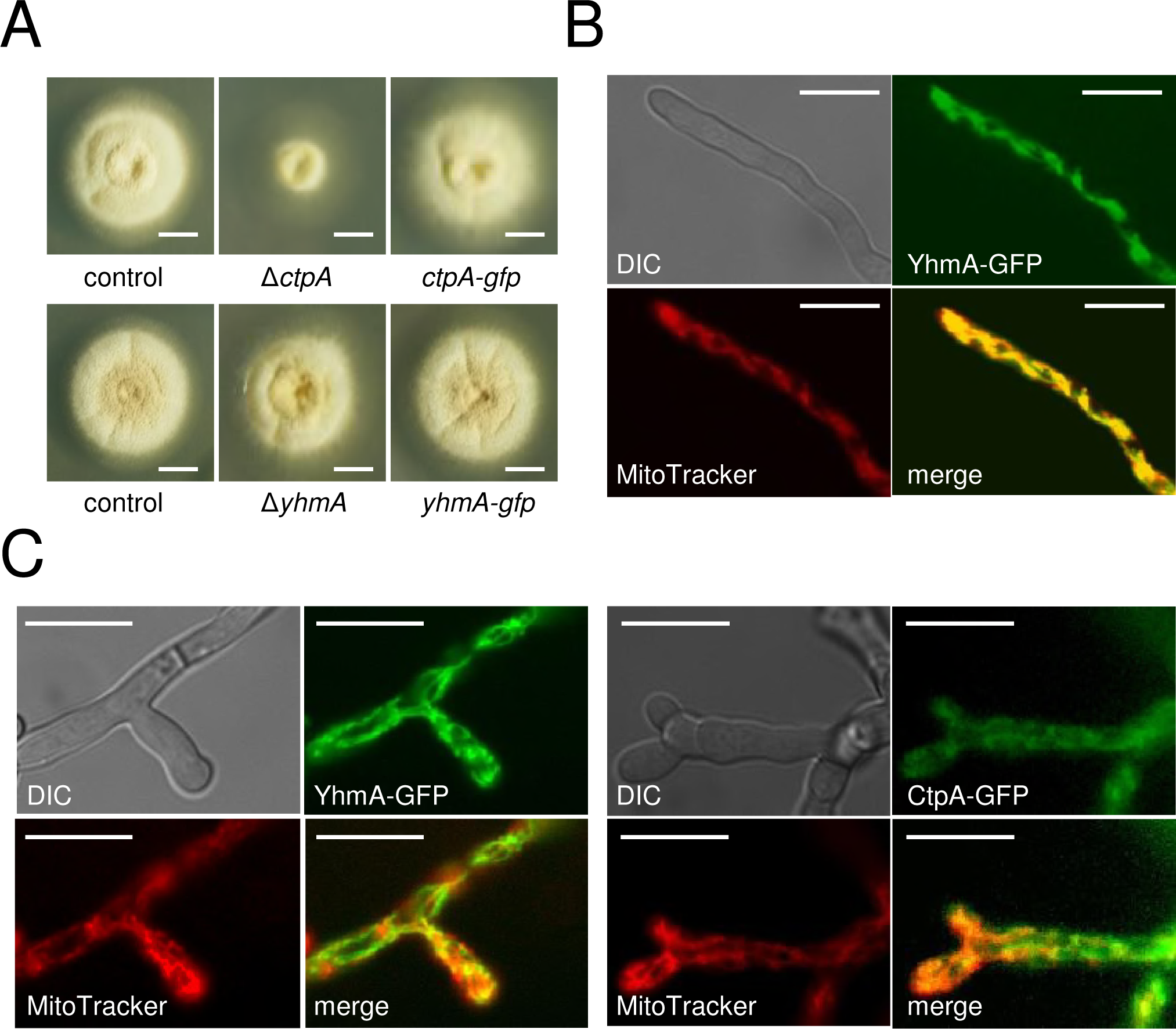
(A) Expression of *ctpA-gfp* and *yhmA-gfp* complement the phenotypes of the *A. kawachii* Δ*ctpA* and Δ*yhmA* strains, respectively. Control, Δ*ctpA*, and *ctpA-gfp* strains were grown on M agar medium at 25° C, whereas the control, Δ*yhmA*, and *yhmA-gfp* strains were grown on M agar medium at 30° C. Scale bars indicate 1 cm. Fluorescence microscopic observation of (B) YhmA-GFP in M medium and (C) in CAP medium and (D) CtpA-GFP in CAP medium. Scale bars indicate 10 μm.

Immunoblot analysis using an anti-GFP antibody indicated that CtpA-GFP and YhmA-GFP were expressed at their predicted molecular weights (59.8 kDa and 61.1 kDa, respectively) (Fig. S4 in the supplemental material). In addition, the bands for both CtpA-GFP and YhmA-GFP exhibited greater intensity with cultivation in CAP medium than M medium, indicating that conditions favorable for citric acid production enhance expression of *ctpA* and *yhmA* at both the mRNA (Fig. 4D) and protein levels.

### Complementation test of *ctpA* and *yhmA* in *S. cerevisiae* strains Δ*ctp1* and Δ*yhm2*

To determine whether *A. kawachii ctpA* and *yhmA* can complement the defect in *S. cerevisiae* strains Δ*ctp1* and Δ*yhm2*, the *ctpA* and *yhmA* genes were expressed in *S. cerevisiae* Δ*ctp1* and Δ*yhm2*, respectively, under control of the respective native promoters.

We first characterized the phenotype of the *S. cerevisiae* Δ*ctp1* strain, because no phenotypic change was observed following disruption of *ctp1* (24). We performed a spot growth assay under cultivation conditions including low temperature stress at 15°C, cell wall stress (Congo red and calcofluor white), and varying carbon source (glucose, acetate, or glycerol). However, as no phenotypic changes were observed following disruption of *ctp1* (data not shown), complementation testing was not possible for *ctpA*.

The *S. cerevisiae* Δ*yhm2* strain reportedly exhibits a growth defect in acetate (SA) medium but not in glucose (SD) medium (Fig. 6) (31). Complementation of *yhmA* (Δ*yhm2* + *yhmA*) remedied the deficient growth of the Δ*yhm2* strain in SA medium as well as the positive control vector carrying *YHM2* (Δ*yhm2* + *YHM2*), indicating that *yhmA* complements the loss of *YHM2* function in *S. cerevisiae*.

**Figure 6.**
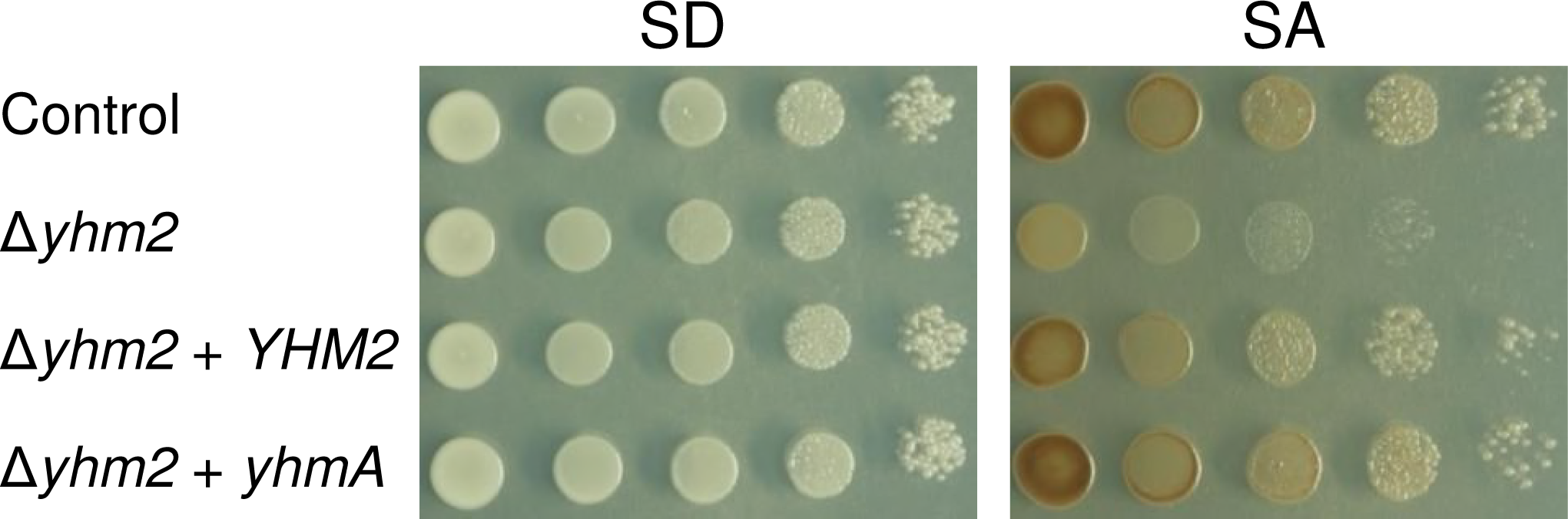
Expression of *yhmA* complements the *S. cerevisiae* Δ*yhm2* phenotype. Ten-fold serial dilutions of 10^7^ cells of control strain, Δ*yhm2*, Δ*yhm2* + *yhm2*, and Δ*yhm2* + *yhmA* cells (all strains pre-cultured for 24 h in SC medium without tryptophan) were inoculated onto SD (glucose) or SA (acetate) medium and incubated at 30° C for 3 days.

### Intracellular levels of nicotinamide cofactors

It was hypothesized that one physiologic role of Yhm2 in *S. cerevisiae* is to increase the NADPH level in the cytosol (31). Thus, we investigated the effect of disrupting *ctpA* and *yhmA* on the intracellular redox state of *A. kawachii*. The control, Δ*ctpA*, and Δ*yhmA* strains were pre-cultivated in M medium at 30°C for 36 h and then transferred to CAP medium and further cultivated at 30°C for 12 h, at which time the NADH/ NAD^+^ ratio, total amount of NADH and NAD^+^, NADPH/NADP^+^ ratio, and total amount of NADPH and NADP^+^ were determined (Table 1).

**Table 1.** Intracellular redox balance of *Aspergillus kawachii* strains.

| Strain | NADH/NAD<sup>+</sup><br>ratio | NADH + NAD<sup>+</sup><br>(μmol/g wet cell) | NADPH/NADP<sup>+</sup><br>ratio | NADPH + NADP<sup>+</sup><br>(μmol/g wet cell) |
| --- | --- | --- | --- | --- |
| Control | 1.07 ± 0.21 | 4.49 ± 0.53 | 1.04 ± 0.0047 | 18.88 ± 0.96 |
| Δ*ctpA* | 0.98 ± 0.20 | 5.31 ± 0.14 | 1.43 ± 0.31 | 18.80 ± 1.62 |
| Δ*yhmA* | 1.17 ± 0.22 | 4.26 ± 0.23 | 0.55 ± 0.17* | 22.64 ± 1.42 |

\* Statistically significant difference ( $p < 0.05$ , Welch's t-test) relative to the result of control strain.

No significant changes were observed with respect to the intracellular NADH/NAD^+^ ratio and total amount of NADH and NAD^+^ in the Δ*ctpA* and Δ*yhmA* strains compared with the control strain. However, the intracellular NADPH/NADP^+^ ratio in the Δ*yhmA* strain was significantly lower than that in the control strain, though there was no significant difference in the total amount of NADPH and NADP^+^ between the Δ*yhmA* strain and the control strain, indicating that the NADPH level was reduced in the Δ*yhmA* strain compared with the control strain. By contrast, the NADPH/NADP^+^ ratio and total amount of NADPH and NADP^+^ did not significantly differ between the Δ*ctpA* strain and the control strain. This result was consistent with a previous report indicating a reduced NADPH/NADP^+^ ratio in the cytosol of the *S. cerevisiae* Δ*yhm2* strain but not in the cytosol of the Δ*ctp1* strain (31).

### Intracellular amino acid levels

Citric acid cycle intermediates are known to serve as substrates for amino acid synthesis in eukaryotic cells (39). Thus, we investigated whether disruption of *ctpA* and *yhmA* affects the intracellular amino acid levels. To compare the intracellular concentrations of amino acids, *A. kawachii* control, Δ*ctpA*, Δ*yhmA*, and Ptet-*ctpA-S* Δ*yhmA* strains were pre-cultivated in M medium at 30°C for 36 h and then transferred to CAP medium and further cultivated at 30°C for 48 h, at which time amino acid levels in the intracellular fraction were determined.

The intracellular concentration of lysine was significantly lower (0.31- and 0.41-fold) in the Δ*ctpA* and Δ*yhmA* strains, respectively, compared with the control strain (Table 2). Furthermore, the Ptet-*ctpA-S* Δ*yhmA* strain exhibited decreased concentrations of versatile amino acids, including aspartic acid, tyrosine, valine, glutamic acid, glycine, histidine, lysine, and alanine (0.43-, 0.19-, 0.32-, 0.43-, 0.35-, 0.51-, 0.22-, and 0.25-fold reduction, respectively).

**Table 2.**
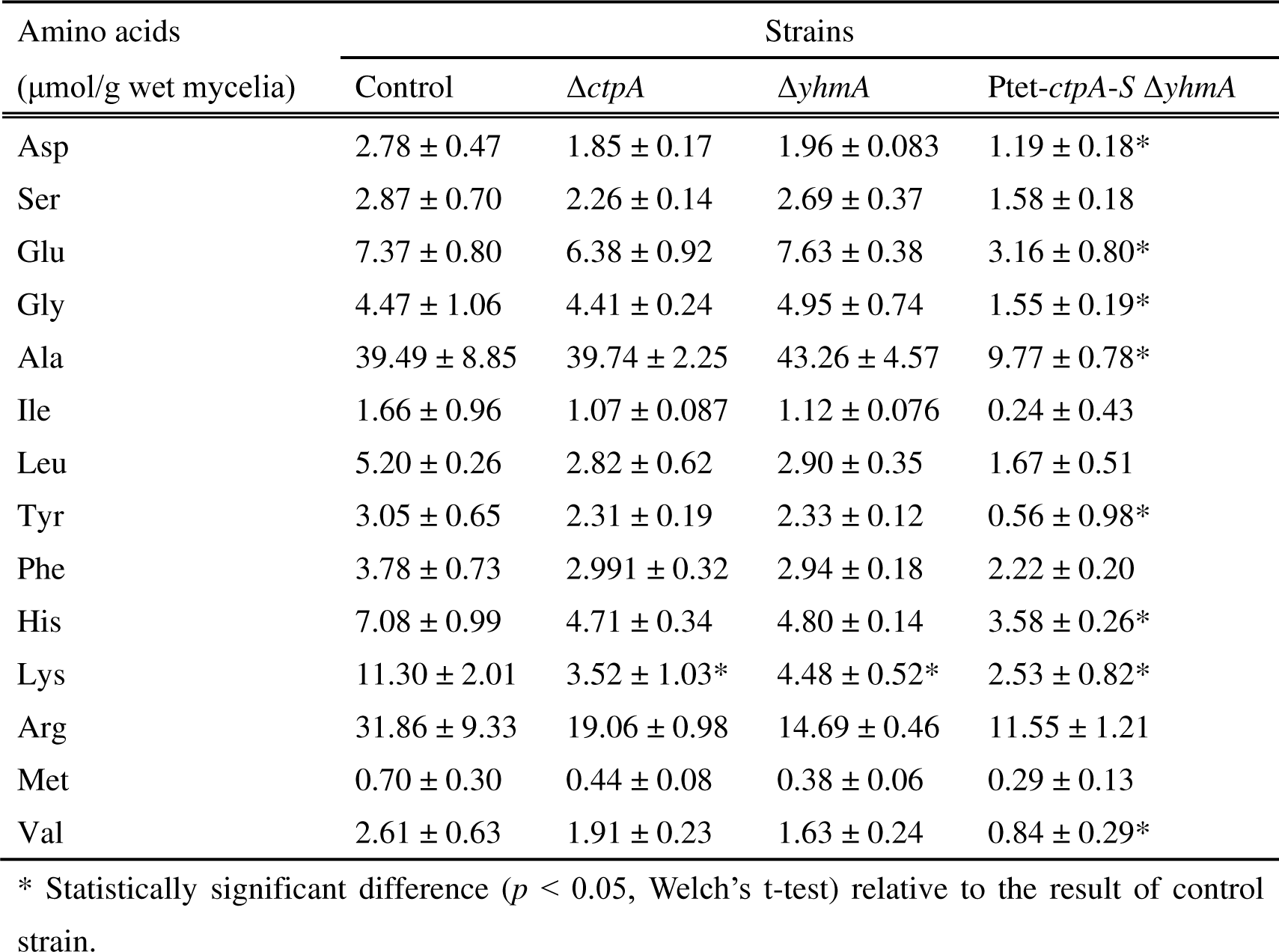
Intracellular amino acid concentrations of *Aspergillus kawachii* strains.

| Amino acids<br>($\mu\text{mol/g}$ wet mycelia) | Strains | | | |
| --- | --- | --- | --- | --- |
| | Control | $\Delta\text{ctpA}$ | $\Delta\text{yhmA}$ | Ptet-*ctpA-S* $\Delta\text{yhmA}$ |
| Asp | $2.78 \pm 0.47$ | $1.85 \pm 0.17$ | $1.96 \pm 0.083$ | $1.19 \pm 0.18^*$ |
| Ser | $2.87 \pm 0.70$ | $2.26 \pm 0.14$ | $2.69 \pm 0.37$ | $1.58 \pm 0.18$ |
| Glu | $7.37 \pm 0.80$ | $6.38 \pm 0.92$ | $7.63 \pm 0.38$ | $3.16 \pm 0.80^*$ |
| Gly | $4.47 \pm 1.06$ | $4.41 \pm 0.24$ | $4.95 \pm 0.74$ | $1.55 \pm 0.19^*$ |
| Ala | $39.49 \pm 8.85$ | $39.74 \pm 2.25$ | $43.26 \pm 4.57$ | $9.77 \pm 0.78^*$ |
| Ile | $1.66 \pm 0.96$ | $1.07 \pm 0.087$ | $1.12 \pm 0.076$ | $0.24 \pm 0.43$ |
| Leu | $5.20 \pm 0.26$ | $2.82 \pm 0.62$ | $2.90 \pm 0.35$ | $1.67 \pm 0.51$ |
| Tyr | $3.05 \pm 0.65$ | $2.31 \pm 0.19$ | $2.33 \pm 0.12$ | $0.56 \pm 0.98^*$ |
| Phe | $3.78 \pm 0.73$ | $2.991 \pm 0.32$ | $2.94 \pm 0.18$ | $2.22 \pm 0.20$ |
| His | $7.08 \pm 0.99$ | $4.71 \pm 0.34$ | $4.80 \pm 0.14$ | $3.58 \pm 0.26^*$ |
| Lys | $11.30 \pm 2.01$ | $3.52 \pm 1.03^*$ | $4.48 \pm 0.52^*$ | $2.53 \pm 0.82^*$ |
| Arg | $31.86 \pm 9.33$ | $19.06 \pm 0.98$ | $14.69 \pm 0.46$ | $11.55 \pm 1.21$ |
| Met | $0.70 \pm 0.30$ | $0.44 \pm 0.08$ | $0.38 \pm 0.06$ | $0.29 \pm 0.13$ |
| Val | $2.61 \pm 0.63$ | $1.91 \pm 0.23$ | $1.63 \pm 0.24$ | $0.84 \pm 0.29^*$ |

\* Statistically significant difference ( $p < 0.05$ , Welch's t-test) relative to the result of control strain.

### Effect of amino acids on growth of the Ptet-*ctpA*-*S* Δ*yhmA* strain

Because we found that disruption of *ctpA* and *yhmA* significantly reduced the concentrations of intracellular amino acids (Table 2), we investigated whether the severe growth defect of the Ptet-*ctpA*-*S* Δ*yhmA* strain in M medium without Dox (Fig. 2C) is due to a defect in amino acid synthesis. The Ptet-*ctpA-S* Δ*yhmA* strain was cultivated in M medium with or without Dox or with various amino acids at a concentration of 0.5% (wt/vol) (Fig. 7A). The defective growth of the Ptet-*ctpA-S* Δ*yhmA* strain was not remedied by supplementation with proline or histidine, but the defect was remedied to some extent by supplementation with aspartic acid, phenylalanine, arginine, and glutamic acid and significantly remedied by supplementation with lysine. Thus, we further examined the effect of lysine at concentrations ranging from 0.2 to 27 mM (27 mM corresponds to 0.5% [wt/vol]). The results confirmed that addition of lysine remedies the growth defect of the Ptet-*ctpA-S* Δ*yhmA* strain in M medium without Dox (Fig. 7B). Together with the intracellular amino acid level data, this result indicates that CtpA and YhmA are required for lysine biosynthesis and that a lack of lysine causes a significant growth defect in the Ptet-*ctpA-S* Δ*yhmA* strain.

**Figure 7.**
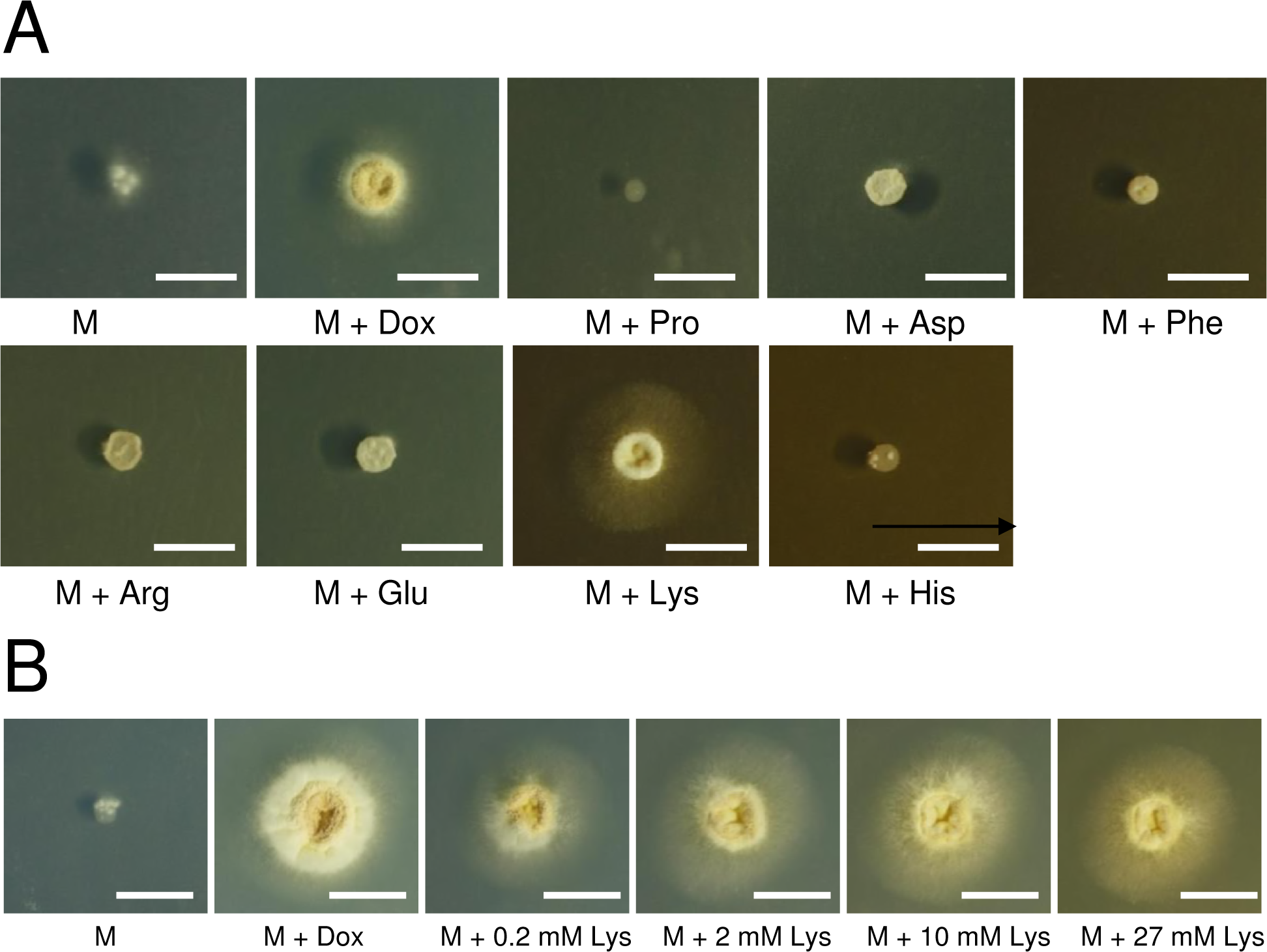
Effect of amino acids on colony formation of the Ptet-*ctpA-S* Δ*yhmA* strain. (A) Conidia (10^4^) of the Ptet-*ctpA-S* Δ*yhmA* strain were inoculated onto M agar medium with or without 1 μg/ml Dox and with 0.5% (wt/vol) various amino acids. (B) Conidia (10^4^) of the Ptet-*ctpA-S* Δ*yhmA* strain were grown on M agar medium with or without 1 μg/ml Dox and with 0.2∼27 mM lysine. The conidia were incubated on the agar medium at 30° C for 4 days. Scale bars indicate 1 cm.

## DISCUSSION

In this study, we attempted to identify the mitochondrial citrate transporters in the citric acid–producing fungus, *A. kawachii*. We identified two candidates, CtpA and YhmA, as mitochondrial citrate transporters in *A. kawachii* based on sequence homology to *S. cerevisiae* Ctp1 and Yhm2, respectively (24, 31). The homologs of Ctp1 are conserved in higher eukaryotes, whereas the homologs of Yhm2 are not conserved in higher eukaryotes such as mammals (31). Interestingly, we found that the *yhmA* gene is conserved downstream of the citrate synthase–encoding gene *citA* in members of the Pezizomycotina, a subphylum of the Ascomycota (Table S1 in the supplemental material). In addition, an RNA-binding protein–encoding gene that is a homolog of *NRD1* in *S. cerevisiae* (40) that localizes upstream of the *citA* gene is also conserved. This gene cluster seems to be conserved in the Pezizomycotina but not in other subdivisions of the Ascomycota, Saccharomycotina (including *S. cerevisiae*), and Taphrinomycotina. Thus, the gene cluster might have arisen during evolution of the Pezizomycotina.

A previous investigation of an *A. niger ctpA* deletion mutant showed that *ctpA* is involved in citric acid production during the early growth stage (28); however, whether this gene was involved in citrate transport remained unclear. Our biochemical experiments confirmed that CtpA and YhmA of *A. kawachii* are citrate transporters. YhmA and CtpA reconstituted proteoliposomes exhibited only counter-exchange transport activity, as previously reported for Ctp1 and Yhm2 (24, 31).

CtpA exhibited citrate transport activity using counter substrates, particularly *cis-*aconitate and malate (Fig. 1A). The substrate specificity of CtpA was very similar to that of yeast Ctp1 and rat CTP, known citrate/malate carriers (18, 37), except that CtpA also exhibited relatively low citrate/citrate exchange activity, unlike Ctp1 and CTP (15, 18, 41). YhmA exhibited citrate transport activity using a wider variety of counter substrates, including citrate, 2-oxoglutarate, malate, *cis*-aconitate, and succinate (Fig. 1B). The substrate specificity of YhmA was also similar to that of Yhm2, with some exceptions (31). Malate and *cis*-aconitate were identified as low-specificity substrates for Yhm2 (31), whereas YhmA exhibited relatively high specificity for malate and *cis*-aconitate.

In analyses of intracellular organic acids, we detected citrate, malate, and 2-oxoglutarate at similar levels of approximately 10 μmol/g wet mycelial weight in the control strain (specifically, citrate: 8.24 μmol/g wet mycelia, malate: 11.07 μmol/g wet mycelia, 2-oxoglutarate: 16.93 μmol/g wet mycelia) (Fig. 3B). Thus, these organic acids appear to be present at comparable concentrations in *A. kawachii* cells. The finding that purified CtpA exhibited higher citrate transport activity when malate was used as the counter substrate, compared with 2-oxoglutarate, suggests that CtpA functions primarily as a citrate/malate carrier *in vivo* (Fig. 8). Purified YhmA exhibited almost equal citrate transport activity when malate or 2-oxoglutarete was used as the counter substrate, suggesting that both malate and 2-oxoglutarate might be physiologic substrates for citrate transport by YhmA.

**Figure 8.**
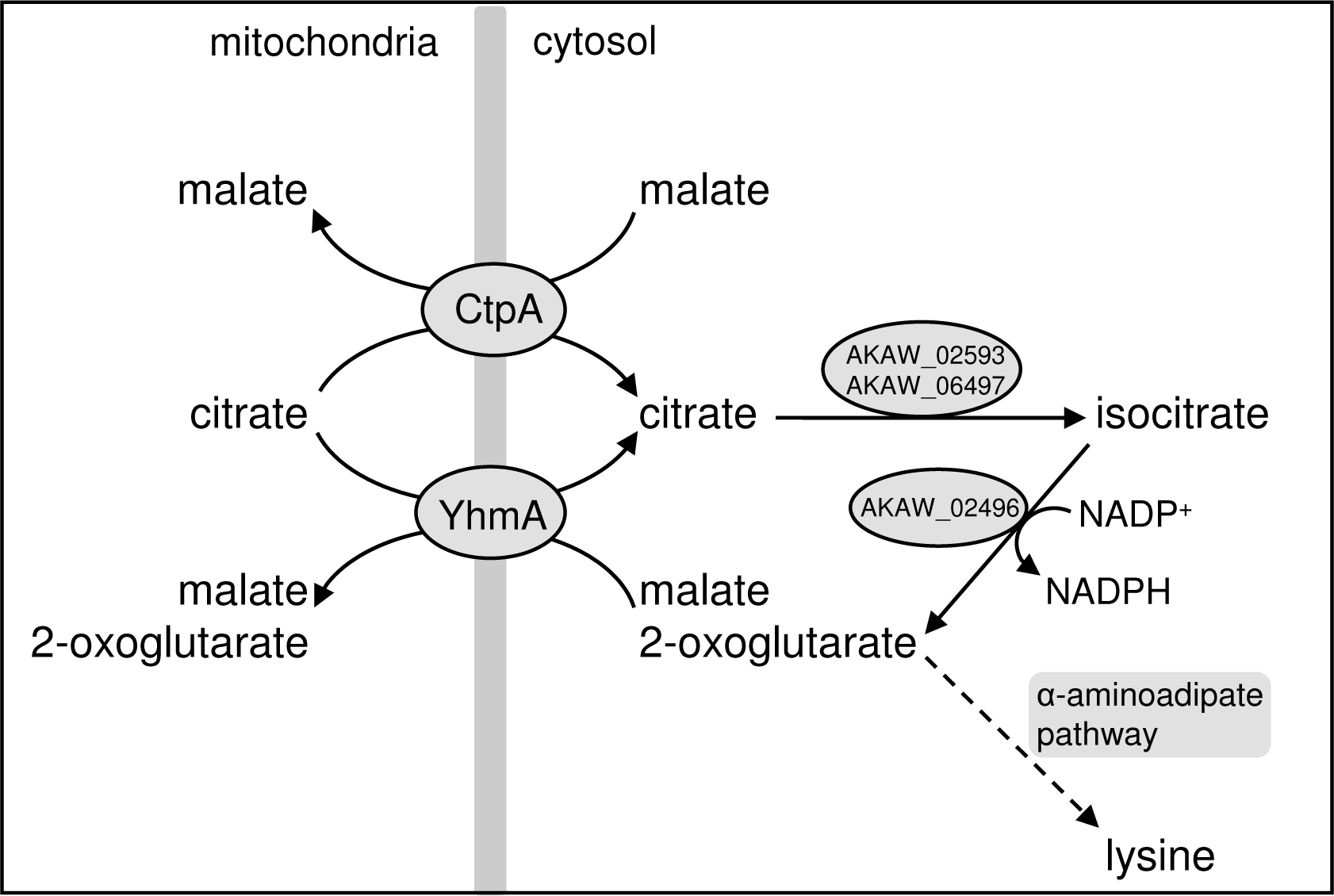
Putative relationships between citrate transport, generation of NADPH, and lysine biosynthesis in *A. kawachii*.

The *A. kawachii yhmA* gene complemented the defective phenotype of the *S. cerevisiae* Δ*yhm2* strain (Fig. 6). Also, disruption of *yhmA* caused a reduction in the NADPH/NADP^+^ ratio, as previously reported in a study of the *S. cerevisiae* Δ*yhm2* strain (Table 1) (31). These results suggest that YhmA plays a role in increasing the NADPH reducing power in the cytosol of *A. kawachii*, similar to *S. cerevisiae* Yhm2 (31). According to metabolic models of *S. cerevisiae* (31) and *A. niger* (42), cytosolic citrate could be converted to 2-oxoglutarate via isocitrate by cytosolic aconitase (AKAW_02593 and AKAW_06497) and NADP^+^-dependent isocitrate dehydrogenase (AKAW_02496) (Fig. 8). During this reaction, NADP^+^ is converted to NADPH by NADP^+^-dependent isocitrate dehydrogenase.

We could not construct a *ctpA* and *yhmA* double disruptant. In addition, downregulation of *ctpA*-*S* in the Ptet-*ctpA-S* Δ*yhmA* strain caused a severe growth defect in M medium (Fig. 2C). These results indicate that double disruption of *ctpA* and *yhmA* causes synthetic lethality in M medium. Disruption of *ctpA* and/or *yhmA* caused a significant reduction in the intracellular lysine concentration (Table 2), and we found that supplementation with lysine relieved the growth defect of the Ptet-*ctpA-S* Δ*yhmA* strain in M medium without Dox (Fig. 7A and B). In fungi, lysine is synthesized from cytosolic 2-oxoglutarate via the α-aminoadipate pathway (43-48). The 2-oxoglutarate available for lysine biosynthesis might be derived primarily from citrate transported by CtpA and YhmA in *A. kawachii* through the metabolic pathway described above (Fig. 8).

The lysine auxotrophic phenotype of the *A. kawachii* Ptet-*ctpA* Δ*yhmA* strain was inconsistent with a previous report indicating that the phenotype of the *S. cerevisiae* Δ*ctp1* Δ*yhm2* strain is very similar to that of the Δ*yhm2* strain (31). Because the prior study used a *lys2-*801 genetic background strain of *S. cerevisiae* (31), the strains were cultivated in medium supplemented with lysine. Thus, we constructed a *ctp1* and *yhm2* double disruptant using *S. cerevisiae* strain W303-1A carrying the *LYS2* gene to clarify whether double disruption of *ctp1* and *yhm2* results in a lysine auxotrophic phenotype in *S. cerevisiae*. However, the Δ*ctp1* Δ*yhm2* strain carrying *LYS2* exhibited a phenotype similar to that of the previously reported Δ*ctp1* Δ*yhm2 lys2-*801 strain (31) (data not shown). Thus, citrate transporters appear to have different physiologic roles in *A. kawachii* and *S. cerevisiae* with respect to lysine biosynthesis. In addition, it was recently reported that the mitochondrial carriers Yhm2, Odc1, and Odc2 are essential for lysine and glutamate biosynthesis in *S. cerevisiae* (49). Because Odc1 and Odc2 function primarily in transporting 2-oxoadipate and 2-oxoglutarate from the mitochondria to the cytosol (50), these 2-oxodicarboxylates are thought to be involved in lysine and glutamate biosynthesis, respectively (49). An Odc1 and Odc2 homolog-encoding gene (AKAW_05597) is present in the genome of *A. kawachii*. Therefore, the physiologic role of this protein in lysine biosynthesis should be further studied to clarify the different roles of mitochondrial transporters in *A. kawachii* and *S. cerevisiae*.

In conclusion, CtpA and YhmA are mitochondrial citrate transporters involved in citric acid production and lysine biosynthesis in *A. kawachii*. *Aspergillus kawachii* is widely used in the shochu fermentation industry in Japan. Thus, our findings are expected to enhance understanding of the citric acid production mechanism and facilitate optimization of strategies to control the activity of *A. kawachii*.

## MATERIALS AND METHODS

### Strains and culture conditions

*Aspergillus kawachii* strain SO2 (51) and *S. cerevisiae* strain W303-1A (52) were used as parental strains in this study (Table S2). Control *A. kawachii* and *S. cerevisiae* strains were defined to show same auxotrophic background for comparison with the respective disruption and complementation strains.

*Aspergillus kawachii* strains were cultivated in M medium (53, Fungal Genetics Stock Center [FGSC] [http://www.fgsc.net/methods/anidmed.html]) with or without 0.211% (wt/vol) arginine and/or 0.15% (wt/vol) methionine or CAP (citric acid production) medium (10% [wt/vol] glucose, 0.3% [wt/vol] (NH_4_)_2_SO_4_, 0.001% [wt/vol] KH_2_PO_4_, 0.05% [wt/vol] MgSO_4_·7H_2_O, 0.000005% [wt/vol] FeSO_4_·7H_2_O, 0.00025% [wt/vol] ZnSO_4_·5H_2_O, 0.00006% [wt/vol] CuSO_4_·5H_2_O [pH 4.0]). CAP medium was adjusted to the required pH with HCl.

*Saccharomyces cerevisiae* strains were grown in YPD medium, synthetic complete (SC) medium, or minimal medium containing 2% (wt/vol) glucose (SD) or 1% (wt/vol) sodium acetate (SA) as a carbon source (54).

### Construction of *ctpA* and *yhmA* disruptants

The *ctpA* and *yhmA* genes were disrupted by insertion of the *argB* gene. The gene disruption cassette encompassing 2 kb of the 5’-end of the target gene, 1.8 kb of *argB*, and 2 kb of the 3’-end of the target gene was constructed by recombinant PCR using the primer pairs AKxxxx-FC/AKxxxx-del-R1, AKxxxx-F2/AKxxxx-R2, and AKxxxx-del-F3/AKxxxx-RC, respectively (where ‘xxxx’ indicates ctpA or yhmA; Table S3 in the supplemental material). For amplification of the *argB* gene, the plasmid pDC1 was used as template DNA (55). The resultant DNA fragment was amplified with the primers AKxxxx-F1 and AKxxxx-R3 and used to transform *A. kawachii* strain SO2, yielding the Δ*ctpA* and Δ*yhmA* strains. Transformants were selected on M agar medium without arginine. Introduction of the *argB* gene into the target locus was confirmed based on PCR using the primer pairs AKxxxx-FC and AKxxxx-RC and the SalI digestion pattern (Fig. S5A and B in the supplemental material). After the confirmation of gene disruption, the Δ*ctpA* and Δ*yhmA* strains were transformed with the *sC* gene cassette to use the same auxotrophic genetic background strains for the comparative study. The *sC* gene cassette was prepared by PCR using *A. kawachii* genomic DNA as template DNA and the primer pair sC-comp-F and sC-comp-R (Table S3 in the supplemental material). Transformants were selected on M agar medium without methionine.

### Construction of complementation strains for the *ctpA* and *yhmA* disruptants

To analyze complementation of the *ctpA* and *yhmA* disruptants with wild type (wt) *ctpA* and *yhmA*, respectively, gene replacement cassettes encompassing 2 kb of the 5’-end of the target gene, 1.4 kb of wt *ctpA* or *yhmA*, 4.2 kb of *sC*, and 1.8 kb of *argB* were constructed by recombinant PCR using the primer pairs AKxxxx-FC/AKxxxx-comp-R1 and AKxxxx-comp-F2/AKxxxx-comp-R2 (where ‘xxxx’ indicates *ctpA* or *yhmA*; Table S3 in the supplemental material). Fragments totaling 6 kb of *sC* and *argB* were simultaneously amplified using a plasmid carrying tandemly connected *sC* and *argB* as the template. Transformants were selected on M agar medium without methionine. Introduction of the wt *ctpA* gene into the *ctpA* disruptant was confirmed by PCR using the primer pairs AKctpA-FC/AKctpA-comp-R2 and AKctpA-comp-F2/AKctpA-comp-R2 (Fig. S5C in the supplemental material). Introduction of the wt *yhmA* gene into the *yhmA* disruptant was confirmed by PCR using the primer pair AKyhmA-FC and AKyhmA-RC (Fig. S5D in the supplemental material).

### Construction of strains expressing CtpA-S and YhmA-S

The pVG2.2 vector (56) was obtained from the FGSC (Manhattan, KS) and used to construct *A. kawachii* strains expressing S-tag–fused CtpA or YhmA under control of the Tet-On promoter. First, the *pyrG* marker gene of pVG2.2 was replaced with the *sC* marker gene. The *sC* gene was amplified by PCR using *A. nidulans* genomic DNA as the template and the primers pVG2.2ANsC-inf-F1 and pVG2.2ANsC-inf-R1 (Table S3 in the supplemental material). The resulting PCR amplicon was cloned into pVG2.2 digested with AscI. Second, the intergenic regions of AKAW_01302 and AKAW_01303 were cloned into the vector for integration into the locus of the *A. kawachii* genome. The intergenic regions of AKAW_01302 and AKAW_01303 were amplified by PCR using *A. kawachii* genomic DNA and the primers pVG2.2ANsC-inf-F2 and pVG2.2ANsC-inf-R2. The resulting PCR amplicons were cloned into the vector digested with PmeI, yielding pVG2.2ANsC.

Next, the *yhmA-S* and *ctpA-S* genes were amplified by PCR using the primer sets pVG2.2ANsC-yhmA-S-inf-F/pVG2.2ANsC-yhmA-S-inf-R and pVG2.2ANsC-ctpA-S-inf-F/pVG2.2ANsC-ctpA-S-inf-R, respectively. The amplified fragments were cloned into the PmeI site of pVG2.2ANsC, yielding pVG2.2ANsC-ctpA-S and pVG2.2ANsC-yhmA-S, respectively. An In-Fusion HD cloning kit (Takara Bio, Shiga, Japan) was used for cloning reactions.

Finally, pVG2.2ANsC-ctpA-S and pVG2.2ANsC-yhmA-S were used to transform the Δ*ctpA* and Δ*yhmA* strains, yielding strains Ptet-*ctpA*-*S* and Ptet-*yhmA*-*S*, respectively. Transformants were selected on M agar medium without methionine. Dox-controlled conditional expression of CtpA-S and YhmA-S was confirmed by immunoblot analysis using anti–S-tag antibody (Medical and Biological Laboratories, Nagoya, Japan) (Figure S3 in the supplemental material).

### Construction of the Ptet-*ctpA-S* Δ*yhmA* strain

To control expression of the *ctpA* gene in the Δ*yhmA* background, we disrupted *yhmA* using the *bar* gene in the Ptet-*ctpA-S* strain. A gene disruption cassette encompassing 2 kb of the 5’-end of the *yhmA* gene, 1.8 kb of *bar*, and 2 kb of the 3’-end of the *yhmA* gene was constructed by recombinant PCR using the primer pairs AKyhmA-FC/AKyhmA-bar-R1, AKymhA-bar-F2/AKyhmA-bar-R2, and AKyhmA-bar-F3/AKyhmA-RC, respectively. For amplification of the *bar* gene, a plasmid carrying *bar* (kindly provided by Prof. Michael J. Hynes, University of Melbourne, Australia) (57) was used as the template DNA. The resultant DNA fragment was amplified with the primers AKyhmA-F1 and AKyhmA-R3 and used to transform *A. kawachii* strain Ptet-*ctpA-S*, yielding the Ptet-*ctpA-S* Δ*yhmA* strain. Transformants were selected on M agar medium with glufosinate extracted from the herbicide Basta (Bayer Crop Science, Bayer Japan, Tokyo, Japan). Introduction of the *bar* gene into the target locus was confirmed by PCR using the primer pair AKyhmA-FC and AKyhmA-RC (Fig. S5E in the supplemental material).

### Construction of strains expressing CtpA-GFP and YhmA-GFP

The plasmid pGS, which carries the *A. kawachii sC* gene (51), was used to construct the expression vector for CtpA-GFP and YhmA-GFP. The genes *ctpA* or *yhmA* (without stop codon) and *gfp* were amplified by PCR using the primer pairs pGS-xxxx-gfp-inf-F1/pGS-xxxx-gfp-inf-R1 and pGS-xxxx-gfp-inf-F2/pGS-gfp-inf-R (where ‘xxxx’ indicates ctpA or yhmA; Table S3 in the supplemental material). For amplification of *gfp*, pFNO3 (58) was used as the template DNA. The amplified fragments were cloned into the SalI site of pGS using an In-Fusion HD cloning kit (Takara Bio).

### Fluorescence microscopy

Strains expressing CtpA-GFP or YhmA-GFP were cultured in M or CAP medium. After cultivation in M medium for 12 h or CAP medium from 14 to 20 h, MitoTracker Red CMXRos (Thermo Fisher Scientific, Waltham, MA) was added to the medium at a concentration of 500 nM and incubated for 40 min. After incubation, the mycelia were washed three times with fresh M or CAP medium and then observed under a DMI6000B inverted-type fluorescent microscope (Leica Microsystems, Wetzlar, Germany). Image contrast was adjusted using LAS AF Lite software, version 2.3.0, build 5131 (Leica Microsystems).

### Construction of the *yhm2* disruptant

The *yhm2* gene was disrupted in *S. cerevisiae* W303-1A by insertion of the *kanMX* gene. The disruption cassette was constructed by PCR using the primer pair SCyhm2-del-F and SCyhm2-del-R, which contained 45 bp of the 5’- and 3’-ends of *yhm2*, respectively (Table S3 in the supplemental material). For amplification of the *kanMX* gene, pUG6 (59) was used as the template DNA. Transformants were selected on YPD agar medium with 200 μg/ml of G418 (Nacalai Tesque, Kyoto, Japan).

### Complementation of *YHM2* and *yhmA* in the *yhm2* disruptant

For the complementation test, we cloned *S. cerevisiae YHM2* and *A. kawachii yhmA* into plasmid YCplac22 carrying *TRP1* (60). Next, 0.6 kb of the 5’-end of *YHM2*, 0.9 kb of *YHM2*, and 0.1 kb of the 3’-end of *YHM2* were amplified by PCR using YCplac22-yhm2-inf-F and YCplac22-yhm2-inf-R. The amplicon was cloned into the SalI site of YCplac22, yielding YCplac22-*yhm2*.

Next, 0.6 kb of the 5’-end of *YHM2* and *yhmA* were amplified by PCR using the primer pairs YCplac22-yhm2-inf-F/YCplac22-yhmA-inf-R1 and YCplac22-yhmA-inf-F2/YCplac22-yhmA-inf-R2, respectively. For amplification of *yhmA* without the intron, *A. kawachii* cDNA was used as the template. The cDNA from *A. kawachii* was prepared using RNAiso Plus (Takara Bio) and reverse-transcription using SuperScript IV (Thermo Fisher Scientific). The amplified fragments were inserted into the SalI site of YCplac22, yielding YCplac22-*YHM2* and YCplac22-*yhmA*, respectively. An In-Fusion HD cloning kit (Takara Bio) was used for the cloning reactions. The resultant plasmids, YCplac22-*YHM2* and YCplac22-*yhmA*, were transformed into the *S. cerevisiae* Δ*yhm2* strain, yielding Δ*yhm2* + *yhm2* and Δ*yhm2* + *yhmA*, respectively. Transformants were selected on SC agar medium without tryptophan.

### Purification of CtpA-S and YhmA-S

A single-step purification method based on S-tag and S-protein affinity (61) was employed for purification of S-tagged CtpA and YhmA from the *A. kawachii* Ptet-*ctpA-S* and Ptet-*yhmA-S* strains, respectively. The Ptet-*ctpA-S* and Ptet*-yhmA-S* strains were cultured in M medium containing 20 μg/ml Dox with shaking (163 rpm) at 30°C for 36 h and then harvested by filtration. The mycelia were ground to a powder using a mortar and pestle in the presence of liquid nitrogen. A total of 1 g wet weight of powdered mycelia was dissolved in 13 ml of ice-cold extraction buffer (25 mM HEPES [pH 6.8], 300 mM NaCl, 0.5% NP-40, 250 μg/ml phenylmethylsulfonyl fluoride [PMSF], cOmplete [EDTA-free protease inhibitor cocktail, Roche, Basel, Switzerland]) and vigorously mixed using a vortexer. Cell debris was removed by centrifugation at 1,000 × *g* at 4°C for 5 min. The resulting supernatant was centrifuged at 18,800 × *g* at 4°C for 15 min. The supernatant was stirred for 2 h at 4°C. Then, S-protein agarose (Merck Millipore, Darmstadt, Germany) was added to the supernatant, and the resulting mixture was gently mixed for 1 h at 4°C using a rotator. S-protein agarose was collected by centrifugation at 500 × *g* for 5 min and then washed once with extraction buffer (containing 0.2% NP-40, 50 μg/ml PMSF), followed by 5 washes using wash buffer (25 mM HEPES [pH 6.8], 300 mM NaCl, 20 μg/ml PMSF, cOmplete [Roche]). CtpA-S and YhmA-S protein was eluted from the S-protein agarose by mixing with elution buffer (25 mM HEPES [pH 6.8], 300 mM NaCl, 0.1% NP-40, 3 M MgCl_2_·7H_2_O) and incubating at 37°C for 10 min. The eluted protein was desalted using Vivacon 500 ultrafiltration units (Sartorius, Gottingen, Germany) with a >10 kDa molecular weight cut-off membrane and washed with buffer (25 mM HEPES [pH 6.8], 300 mM NaCl, 0.1% NP-40) 5 times. The concentrations of CtpA-S and YhmA-S were determined using a Qubit protein assay kit (Thermo Fisher Scientific). The purified proteins were subjected to sodium dodecyl sulfate–polyacrylamide gel electrophoresis (SDS-PAGE) to confirm purity (Fig. S2 in the supplemental material).

### Transporter assay

YhmA-S or CtpA-S reconstituted proteoliposomes in the presence or absence of internal substrate were prepared using a freeze/thaw sonication procedure (62). Briefly, liposomal vesicles were prepared by probe-type sonication using a Sonifier 250A (Branson Ultrasonics, Division of Emerson Japan, Kanagawa, Japan) with 100 mg of L-α-phosphatidylcholine from egg yolk (Nacalai Tesque) in buffer G (10 mM PIPES, 50 mM NaCl, and 1 mM organic acids [oxaloacetate, succinate, cis-aconitate, citrate, 2-oxoglutarate, or malate]). Solubilized CtpA-S and YhmA-S (500 ng of each) was added to 500 μl of liposomes and immediately frozen in liquid nitrogen and then sonicated after melting. Extraliposomal substrate was removed using Bio Spin 6 columns (Bio-Rad, Hercules, CA). To initiate the transport reaction, 1 mM [1,5-^14^C]-citrate (18.5 kBq) (PerkinElmer, Waltham, MA) was added and incubated at 37°C for 30 min. After the reaction, extraliposomal labeled and nonlabeled substrates were removed using Bio Spin 6 columns (Bio-Rad). Intraliposomal radioactivity was then measured using a Tri-Carb 2810TR liquid scintillation analyzer (PerkinElmer) after mixing with Ultima Gold scintillation cocktail (PerkinElmer).

### Measurement of intracellular nicotinamide cofactor levels

To determine the intracellular levels of nicotinamide cofactors, conidia (2 × 10^7^ cells) of *A. kawachii* control, Δ*ctpA*, and Δ*yhmA* strains were inoculated into 100 ml of M medium and pre-cultured with shaking (163 rpm) at 30°C for 36 h. The mycelia were then transferred to 50 ml of CAP medium and further cultivated for 12 h. The mycelia were collected and ground to a fine powder using a mortar and pestle in the presence of liquid nitrogen. Next, 10 ml of cold PBS (1.37 M NaCl, 81 mM Na_2_HPO_4_, 26.8 mM KCl, 14.7 mM KH_2_PO_4_) was added to 1 g of mycelial powder, vortexed, and then centrifuged at 138,000 × *g* at 4°C for 30 min. The NADH/NAD^+^ ratio, total amount of NADH and NAD^+^, NADPH/NADP^+^ ratio, and total amount of NADPH and NADP^+^ were measured using an Amplite Fluorimetric NAD/NADH ratio assay kit (red fluorescence) and Amplite Fluorimetric NADP/NADPH ratio assay kit (red fluorescence) (AAT Bioquest, Sunnyvale, CA), respectively, according to the manufacturer’s protocol. Fluorescence was detected using an Infinite M200 FA (Tecan, Männedorf, Switzerland).

### Measurement of extracellular and intracellular organic acids

To measure levels of extracellular and intracellular organic acids, conidia (2 × 10^7^ cells) of *A. kawachii* control, Δ*ctpA*, Δ*yhmA* strains were inoculated into 100 ml of M medium and pre-cultivated with shaking (180 rpm) at 30°C for 36 h and then transferred to 50 ml of CAP medium and further cultivated with shaking (163 rpm) at 30°C for 48 h. The Ptet-*ctpA*-*S* Δ*yhmA* strain was pre-cultured in M medium with 1 μg/ml Dox and transferred to CAP medium without Dox. The culture supernatant was filtered through a 0.2-μm pore size PTFE filter (Toyo Roshi Kaisha, Japan) and used as the extracellular fraction. Mycelia were used for preparation of the intracellular fraction using a hot water extraction method (63), with modifications. After mycelial growth was measured as wet weight, the mycelia were ground to a powder using a mortar and pestle in the presence of liquid nitrogen. The mycelia were then dissolved in 10 ml of hot water (80°C) per 1 g of mycelial powder, vortexed, and then centrifuged at 138,000 × *g* at 4°C for 30 min. The supernatant was filtered through a 0.2-μm pore size filter and used as the intracellular fraction.

The concentrations of organic acids in the extracellular and intracellular fractions were determined using a Prominence HPLC system (Shimadzu, Kyoto, Japan) equipped with a CDD-10AVP conductivity detector (Shimadzu). The organic acids were separated using tandem Shimadzu Shim-pack SCR-102H columns (300 × 8 mm I.D., Shimadzu) at 50°C using 4 mM *p*-toluenesulfonic acid monohydrate as the mobile phase at a flow rate of 0.8 ml/min. The flow rate of the post-column reaction solution (4 mM *p*-toluenesulfonic acid monohydrate, 16 mM bis-Tris, and 80 μM EDTA) was 0.8 ml/min.

### Measurement of intracellular amino acids

Intracellular fractions of *A. kawachii* strains were prepared as described above. Amino acids were analyzed using a Prominence HPLC system (Shimadzu) equipped with a fluorescence detector (RF-10AXL, Shimadzu) according to a post-column fluorescence derivatization method. Separation of amino acids was achieved using a Shimadzu Shim-pack Amino-Na column (100 × 6.0 mm I.D., Shimadzu) at 60°C and a flow rate of 0.6 ml/min using an amino acid mobile phase kit, Na type (Shimadzu). The fluorescence detector was set to excitation/emission wavelengths of 350/450 nm. The reaction reagents were taken from the amino acid reaction kit (Shimadzu) and maintained at a flow rate of 0.2 ml/min.

### Transcription analysis

For RNA extraction from mycelia, conidia (2 × 10^7^ cells) of *A. kawachii* control strain were inoculated into 100 ml of M medium and cultured for 24, 30, 36, 48, 60, and 72 h at 30°C. For RNA extraction from conidia, conidia (2 × 10^5^) were spread onto M agar medium and cultivated at 30°C for 5 days. After incubation, mycelia and conidia were collected and ground to a powder in the presence of liquid nitrogen. RNA was extracted using RNAiso Plus (Takara Bio) according to the manufacturer’s protocol and then quantified using a NonoDrop-8000 (Thermo Fisher Scientific). cDNA was synthesized from total RNA using a PrimeScript Perfect real-time reagent kit (Takara Bio) according to the manufacturer’s protocol. Real-time RT-PCR was performed using a Thermal Cycler Dice real-time system MRQ (Takara Bio) with SYBR Premix Ex Taq II (Tli RNaseH Plus) (Takara Bio). The following primer sets were used: AKyhmA-RT-F and AKyhmA-RT-R for *yhmA*, AKctpA-RT-F and AKctpA-RT-R for *ctpA*, AKwetA-RT-F and AKwetA-RT-R for *wetA*, and AKactA-RT-F and AKactA-RT-R for *actA* (Table S3 in the supplemental material).

## ACKNOWLEDGMENTS

This study is supported in part by Yonemori Seishin Ikuseikai, Sasakawa Scientific Research Grant from the Japan Science Society, Institute for Fermentation, Osaka (IFO), and by a Grant-in-Aid for Scientific Research (C) (no. 16K07672). C. K. was supported by a Grant-in-Aid for JSPS Research Fellows (no. 17J02753).

